# Comparison of the knockdown resistance locus (*kdr*) in *Anopheles stephensi*, *An. arabiensis*, and *Culex pipiens s.l.* suggests differing mechanisms of pyrethroid resistance in east Ethiopia

**DOI:** 10.1101/2020.05.13.093898

**Authors:** Tamar E. Carter, Araya Gebresilassie, Shantoy Hansel, Lambodhar Damodaran, Callum Montgomery, Victoria Bonnell, Karen Lopez, Daniel Janies, Solomon Yared

## Abstract

The malaria vector, *Anopheles stephensi*, which is typically restricted to South Asia and the Middle East, was recently detected in the Horn of Africa. Controlling the spread of this vector could involve integrated vector control that considers the status of insecticide resistance of multiple vector species in the region. Previous reports indicate that the knockdown resistance mutations (*kdr*) in the voltage-gated sodium channel (*vgsc*) are absent in both pyrethroid resistant and sensitive variants of *An. stephensi* in east Ethiopia but similar information on other vector species in the same areas is limited. In this study, *kdr* and the neighboring intron was analyzed in *An. stephensi*, *An. arabiensis*, and *Culex pipiens s. l*. collected in east Ethiopia between 2016 and 2017. Sequence analysis revealed that all of *Cx. pipiens s.l.* (n = 42) and 71.6% of the *An. arabiensis* (n=67) carried *kdr* L1014F known to confer target-site pyrethroid resistance. Intronic variation was only observed in *An. stephensi* (segregating sites = 6, haplotypes = 3) previously shown to have no kdr mutations. In addition, no evidence of non-neutral evolutionary processes was detected at the *An. stephensi kdr* intron which further supports target-site mechanism not being a major resistance mechanism in this *An. stephensi* population. Overall, these results suggest differences in evolved mechanisms of pyrethroid/DDT resistance in populations of vector species from the same region. Variation in insecticide resistance mechanisms in East Ethiopian mosquito vectors highlight possible species or population specific biological factors and distinct environmental exposures that shape their evolution.

## BACKGROUND

Vector-borne diseases are major public health concern, of which malaria remains a leading threat with 228 million cases reported in 2018 1. In Ethiopia, where both *Plasmodium vivax* and *P. falciparum* are prevalent and multiple *Anopheles* vector populations are present,1.5 million malaria cases were reported in 2017 2. Malaria control in Ethiopia and the rest of Africa is now challenged with the recent discoveries of *An. stephensi*, a malaria vector, which is typically restricted to South Asia and the Middle East, in the Horn of Africa and recently demonstrated to transmit local *Plasmodium* 3, 4, 5, 6. Among several approaches to mitigating the *An. stephensi* is integrated vector control that target multiple vectors. Integrated vector control has the benefits of cutting costs and while minimizing adverse outcomes of single-target vector control on non-target species populations. 7

Integrated vector control strategies based on insecticides should account for insecticide resistance status of the different vectors. In Ethiopia, insecticides like pyrethroids have been deployed through indoor residual spraying and long-lasting insecticidal nets (LLIN). Exacerbated by the use of insecticides in the agricultural industries, widespread insecticide resistance has been reported across multiple vector species 8. In Culicidae, the main mechanisms of resistance to pyrethroids include target-site and metabolic-based resistance 9. Pyrethroid based target-site resistance is caused by mutations in the voltage-gated sodium channel leading to altered neurological response to insecticides in mosquitoes [i.e. knockdown resistance (*kdr*), reviewed in 10]. Knockdown resistance is broadly studied and is widely reported across species of Culicidae including *Anopheles* spp. 9 and *Culex pipiens s.l*. 11. In *Anopheles*, *kdr* involves the substitution of leucine (TTA) with phenylalanine (TTT) or serine (TCA) in the voltage gated sodium channel protein, commonly known as *kdr* mutations L1014F and L1014S 12. Similar mutations that confer resistance to pyrethroids (also known as L1014F and L1014S) are observed in the *vgsc* of *Culex* mosquitoes.

For metabolic resistance, the insecticide is degraded, sequestered or exported out of the cell before it can bind to its target 9. Metabolic resistance has not been linked to a single trackable genetic variant in most species. However, previous functional studies have found the over-expression of detoxification enzymes such as cytochrome P450s lead to metabolic resistance 9, 13.

In Ethiopia, pyrethroid and DDT resistance have been reported in much of the northern and western portion of the country in the primary malaria vector *An. arabiensis* 14, 15, 16, 17. In *An. arabiensis*, both target-site and metabolic resistance play a role in pyrethroid and DDT resistance. In eastern Ethiopia, a recent investigation revealed *An. stephensi* were resistant to pyrethroids, although, the L1014F and L1014S mutations were absent 18. *An. arabiensis* insecticide resistance in eastern Ethiopia has not been well characterized. Even more so, the status of insecticide resistance in *Cx. pipiens s.l.* (most likely *Cx. quinquefasciatus*) is unknown throughout most of the country.

Knowing the status of resistance to pyrethroids across vector species in a region can provide insight into the effectiveness of particular insecticides used to target multiple species. Genetic analyses of putative insecticide resistance loci across local vector populations, can provide information on the range of mechanisms of insecticide resistance in a region. While *kdr* L1014F and L1014S mutation frequencies provide preliminary evidence of target-site resistance to pyrethroids, analysis of the variation in neighboring intronic region provides information of the long-term impact of pyrethroids on the evolution of the mosquito populations. Tests for neutrality, such as Tajima’s D 19, can be used to evaluate the genetic diversity of the kdr locus including the intronic region to determine if the patterns differ from expectations under neutral evolution. It is expected that if the *kdr* locus was under selection due to pressure from the pyrethroids, then we hypothesize that a selective sweep would have led to decreased nucleotide diversity of linked alleles 20, 21. Thus, these analyses are helpful in clarifying the mechanisms of resistance, the current status of pyrethroid resistance, and predicting the risk of resistance emerging locally. Here we examine the nucleotide diversity surrounding the *kdr* locus to test for the hypothesis of selective sweeps in *An. stephensi, An. arabiensis*, and *Culex pipiens s. l.* collected in east Ethiopia.

## METHODS

The study involved sequencing of a portion of the *vgsc* gene that contains loci that when mutated can confer resistance to pyrethroids. For *An. stephensi*, data came from sequences generated in a previous study 18 and generated in the present study. *An. arabiensis* and *Culex* sequence data was also generated in this study as detailed below.

### Sample collection and species identification

*An. stephensi* were collected from Kebri Dehar in 2016 as part the first detection of this species in Ethiopia 4. Mosquitoes were collected as larvae and lab-reared for testing for resistance to insecticides as previously detailed 18. *An. arabiensis* and *Culex* specimens collected in east Ethiopia in 2017 were included in this study. *An. arabiensis* species identification was based on morphological keys and molecular analysis of internal transcribed spacer 2 (*ITS2*) and cytochrome oxidase I (*COI*) loci as reported previously 22. *An. arabiensis* were collected using CDC light traps (John W. Hock, Gainesville, FL, USA) over four different collection times at two sites, Meki (east-central Ethiopia) and Harewe (northeast) in 2017. Harewe and Meki are about 350 km northwest and 600 km west of Kebri Dehar, respectively (Fig 1).

**Fig. 1.**
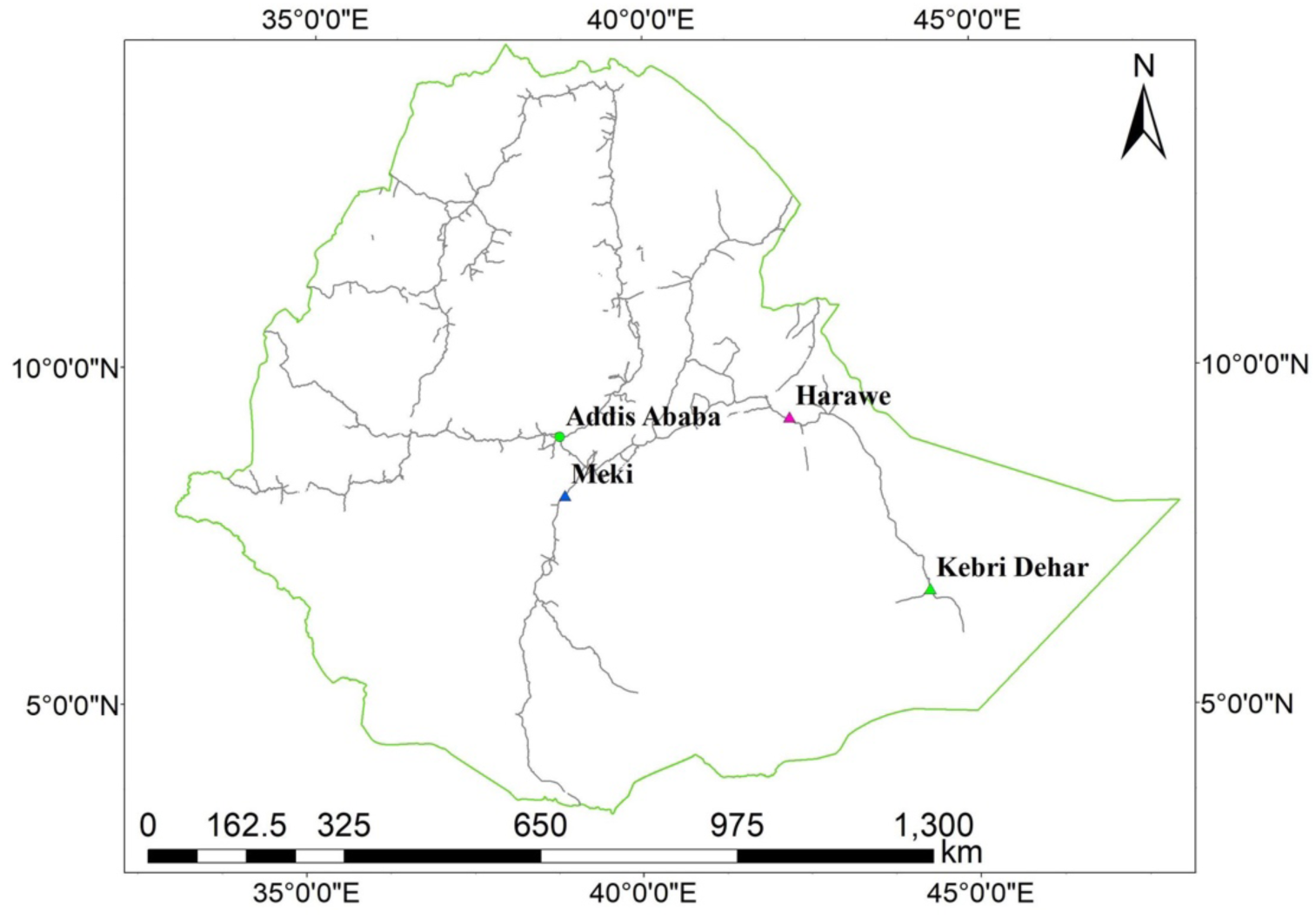
Collection sites.

*Culex* specimens were collected using CDC light traps in Kebri Dehar in 2017. Morphological key and sequencing of *ITS2* locus were used for *Culex* identification using a previously published PCR protocol 4. All amplicons were cleaned using Exosap and sequenced using Sanger technology with ABI BigDyeTM Terminator v3.1 chemistry (Thermofisher, Santa Clara, CA) according to manufacturer recommendations and run on a 3130 Genetic Analyzer (Thermo Fisher, Santa Clara, CA). Sequences were cleaned and analyzed using CodonCode Aligner Program V. 6.0.2 (CodonCode Corporation, Centerville, MA). ITS2 sequences from *Culex* specimen were submitted as queries to the National Center for Biotechnology Information’s (NCBI) Basic Local Alignment Search Tool (BLAST) for species identification 23.

### Amplification and sequencing of kdr loci

Once species or species complex identification was complete, samples were processed. For *kdr* mutation analysis, polymerase chain reaction (PCR) was used to amplify the region of the *vgsc* gene that housed the homologous *kdr* 1014 and a neighboring downstream intron in all specimens (reference sequences used for *An. stephensi*, *An. arabiensis* and *Culex pipiens sl*. were JF304952, GU248311, and BN001092, respectively). One leg from each mosquito specimen or extracted DNA was used as individual templates for PCR. Each species required a different PCR protocol. DNA extraction were performed using DNEasy Qiagen kit (Qiagen, Valencia, USA). All PCR reactions were performed at 25µl total with 12.5 ul 2X Promega Hot Start Master Mix (Promega Corporation, Madison, USA) and the primer conditions listed in Tab 1. *An. stephensi kdr* amplification was completed according to Singh et al., 24 with modifications as detailed in Yared et al., 18. Temperature cycling was as follows: 95°C for 5 min, followed by 35 cycles of 95°C for 30 sec, 50°C for 30 sec, 72°C for 45 sec, and a final extension of 72 °C for 7 min. Amplifications of the kdr fragment from *An. arabiensis* were completed according to methods in Verhaeghen et al 25. Temperature cycling was as follows: 95°C for 1 min, followed by 30 cycles of 95°C for 30 sec, 52°C for 30 sec, 72°C for 1 min, and a final extension of 72°C for 10 min. Amplifications of the kdr fragment from *Culex pipiens* s.*l* were completed according methods in Chen et al 26. Temperature cycling was as follows: 94°C for 5 min, followed by 30 cycles of 94°C for 40 sec, 58°C for 30 sec, 72°C for 40 sec, and a final extension of 72°C for 8 min.

**Tab 1.**
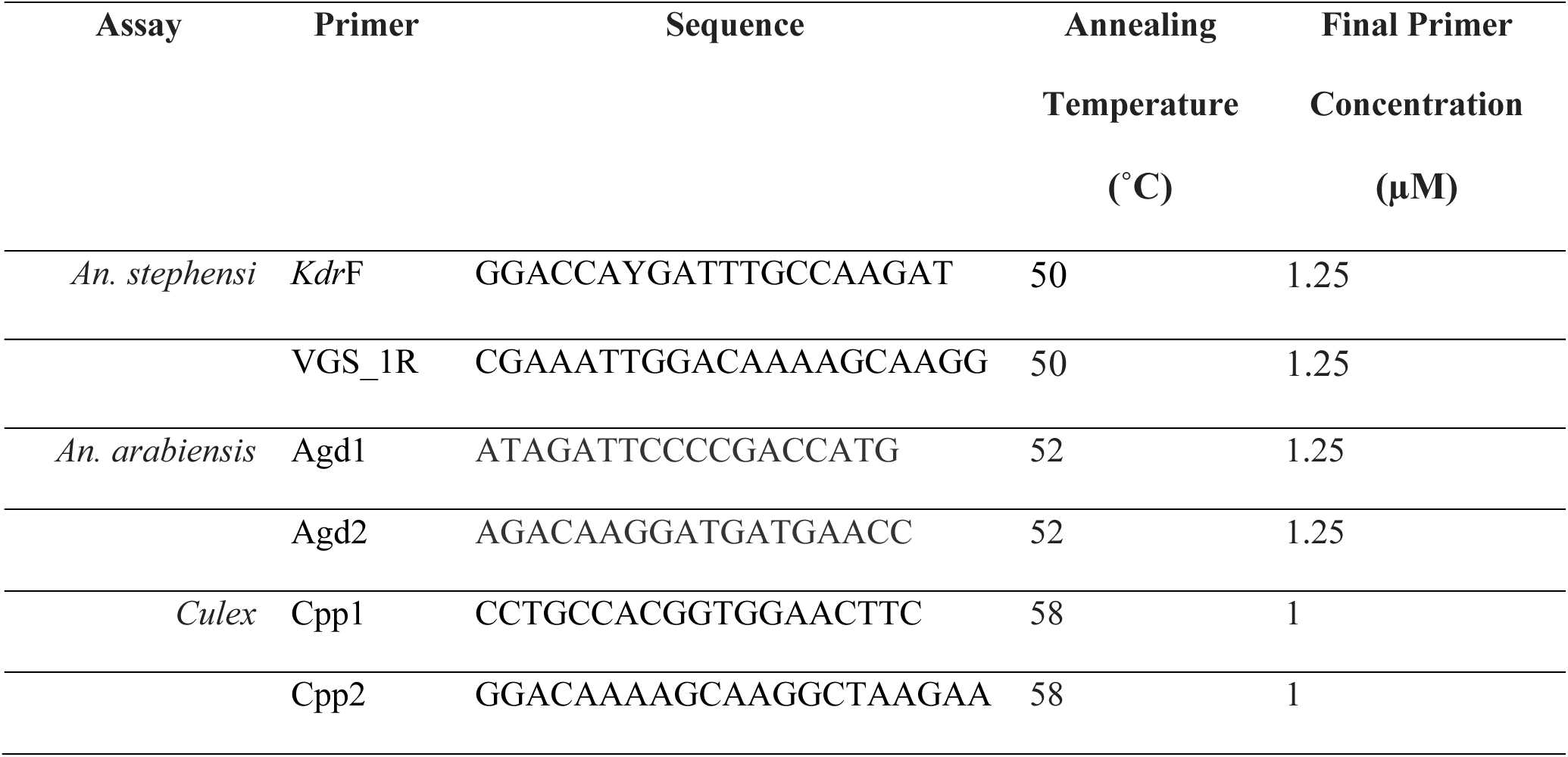
List of primer and conditions used for PCR amplification of portions of the voltage gated sodium channel gene.

All amplicons were cleaned using Exosap and sequenced using Sanger technology with ABI BigDyeTM Terminator v3.1 chemistry (Thermofisher, Santa Clara, CA) according to manufacturer recommendations and run on a 3130 Genetic Analyzer (Thermo Fisher, Santa Clara, CA).

### Sequence analysis

Sequences were submitted as queries to the National Center for Biotechnology Information’s (NCBI) Basic Local Alignment Search Tool (BLAST) to confirm correct loci were amplified. Sequences were then aligned in CodonCode (CodonCode Corp., Dedham, MA, USA) by species or species complex to identify *kdr* L1014F or L1014S mutations based on reference sequence details from previous reports 18, 24, 25. Heterozygous genotypes at kdr was determined based on the number of peaks observed in the chromatogram, with each peak indicate a different alleles. The *kdr* allele and genotype frequencies were then calculated and compared across species.

We determined the level of diversity in the neighboring intron downstream of the *kdr* 1014 in *Culex* spp., *An. arabiensis, and An. stephensi* for additional evidence of selection on that locus. In addition to the sequences generated in this study, we included sequences from resistant and non-resistant *An. stephensi* analyzed in a previous study on insecticide resistance in *An. stephensi* 18. We calculated the number of segregating sites, nucleotide diversity, the estimated number of haplotypes, and haplotype diversity using the program DNAsp v5 27. Haplotypes were reconstructed using Phase 2.1 28, HAPAR, and fastPHASE 29 algorithms in DNAsp. The neighboring downstream intron was also tested for neutrality using Tajima’s D 19, Fu’s F 30, and Fu and Li’s D* and F* tests 31.

## RESULTS

Prior to insecticide resistance genotyping, all *Culex* ITS2 sequences were analyzed to identify species. All sequences were identical and had equivalent high matching scores for two members of the *Cx. pipiens* complex: *Cx. p. quinquefasciatus* and *Cx. p. pipiens*. Because we could not identify these mosquitos to species, we will refer to these specimens by the broader taxonomic classification, *Cx. pipiens s. l.* (i.e., *Cx. pipiens* complex) in this study. *An. arabiensis* species identification was detailed in previous study 22. In total, 10, 33, and 24 *An. arabiensis* were collected in Harewe in November 2016, Harewe in July/August 2017, and Meki in July 2017 collections, respectively.

### Kdr analysis

The *kdr* fragments were sequenced for *An. stephensi*, *Cx. pipiens s.l.*, and *An. arabiensis.* The sequencing resulted in 184, 452, and 290 base pair fragments for *An. stephensi*, *Cx. pipiens s. l*. and *An. arabiensis*, respectively. The percent of each *kdr* genotype observed by species is shown in Fig 2. A total of 131 *An. stephensi* were analyzed, including 80 newly reported sequences. None of the *An. stephensi* analyzed in this study carried a mutation at the *kdr* 1014. All 42 *Cx. pipiens s.l*. specimens collected at the same site carried *kdr* L1014F mutations as homozygous. Of the 67 *An. arabiensis*, 71.6% carried the *kdr* L1014F mutation (heterozygous and homozygous). The allele frequency of L1014F mutation varied across *An. arabiensis* collections, where the highest frequency was observed in Harewe in November 2016 (100%). L1014F allele frequency for Harewe July/August 2017 and in Meki July 2017 collections were 86.4% and 10%, respectively. No L101S mutations were detected in *Cx. pipiens* s.l. or *An. arabiensis*.

**Fig. 2.**
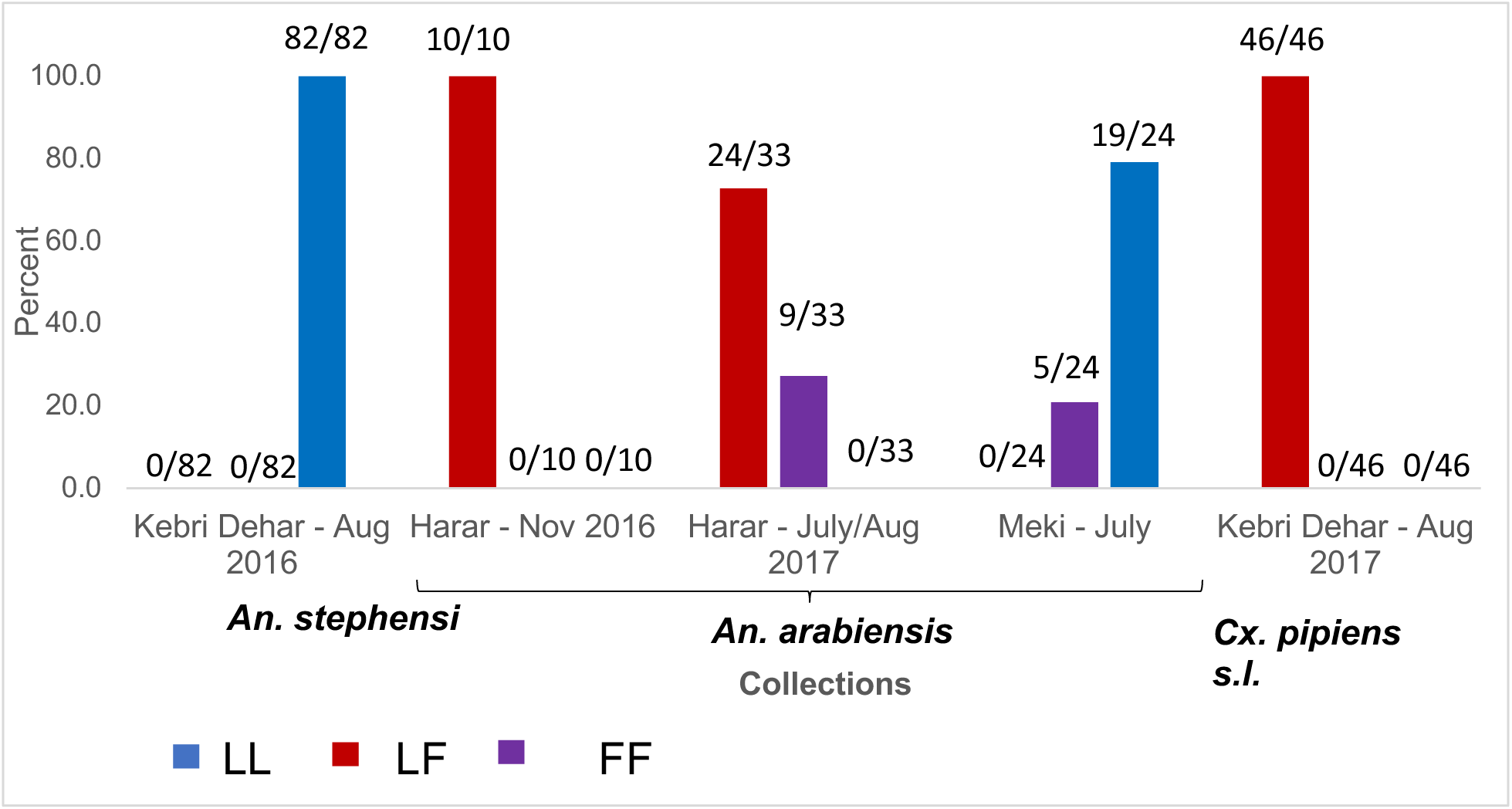
Frequency of *kdr* 1014 genotypes in *An. stephensi, Culex pipiens s.l.*, and *An. arabiensis* collections.

A portion of the neighboring downstream intron for each species was analyzed to evaluate the level of diversity (Fig 3). Intron analysis revealed no polymorphisms for either *Cx. pipiens* or *An. arabiensis* (for both L1014F and L1014 wild type specimens). Of the 131 *An. stephensi* specimens from Kebri Dehar examined for *kdr* mutations, six segregating sites were detected, and three haplotypes predicted. Genetic diversity estimates are reported in Tab 2.

**Fig. 3.**
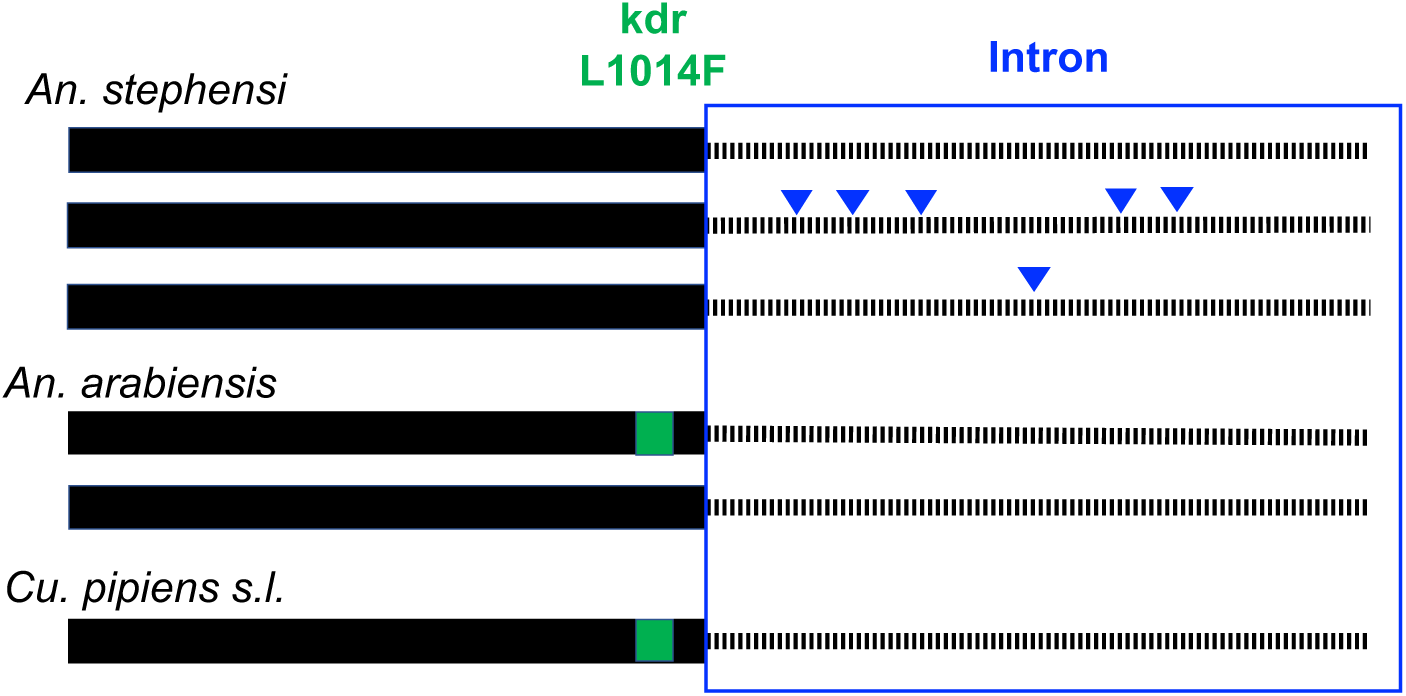
Summary of *kdr* haplotypes across three Culicidae species in east Ethiopia. Solid lines depict the exon housing the *kdr* locus and dotted lines depict the downstream intron. Green square indicates the presence of the *kdr* L1014F. Triangles denote single nucleotide polymorphisms (SNPs) found in the intron relative to the most prevalent intron haplotype.

**Tab 2.** Genetic diversity estimates for *kdr* neighboring downstream intron in the *vgsc* for *An. stephensi*, *An. arabiensis*, and *Cx. pipiens s.l.*, where n = number of genes (two per individuals), S = number of polymorphic (i.e., segregating) sites, K = average number of pairwise nucleotide differences, Pi = nucleotide diversity, h = number of Haplotypes, Hd = haplotype diversity.

| Species | n | S | k | Pi | h | Hd |
| --- | --- | --- | --- | --- | --- | --- |
| *An. stephensi* | 262 | 6 | 0.996 | 0.00545 | 3 | 0.225 |
| *An. arabiensis* | 134 | 0 | 0 | 0 | 1 | 0 |
| *Cx. pipiens s.l.* | 84 | 0 | 0 | 0 | 1 | 0 |

To further evaluate the potential functional significance of the *kdr* locus in *An. stephensi* based on evidence of positive selection, we performed tests for neutrality at the *An. stephensi kdr* intron. No evidence of non-neutral processes was detected in *An. stephensi* for the *kdr* locus (Tab 3). The absence of variation in *An. arabiensis* and *Cx. pipiens* s.l. *kdr* introns precluded tests for neutrality.

**Tab 3.** Tests for neutrality for downstream *kdr* intron for *An. stephensi*. All p-value > 0.10.

| Test | Estimate |
| --- | --- |
| n | 258 |
| Tajima's D | 0.03839 |
| Fu's F | 3.556 |
| Fu and Li's D | 1.04354 |
| Fu and Li's F | 0.82943 |

## DISCUSSION

Our results reveal variation at the *kdr* locus across different vector species found in east Ethiopia suggesting different mechanisms of pyrethroid/DDT resistance. Notably, the *kdr* L1014F mutation was not observed consistently across the species included in this study. Unlike the *An. stephensi*, that carried no L1014F mutations, both *Cx. pipiens sl.* and *An. arabiensis* carried the L1014F. Based on these findings, it is likely that *Cx. pipiens s.l.* and *An. arabiensis* should not share the same mechanisms of pyrethroid resistance as *An. stephensi*. We also observed differences in the nucleotide diversity of the neighboring intronic region of the three species. While *An. stephensi* exhibited multiple segregating sites and resultant haplotypes, only a single intronic haplotype is observed for *An. arabiensis* and *Cx. pipiens s.l.* These data may point to distinct differences in biological and environmental factors that shape each species/population. From a species standpoint, behaviors shaped by both their biology and environment, like feeding and resting preferences may impact the degree of exposure to insecticides. *Cx. pipiens sl* tend toward exophilic behavior and feed generally during the day up to the early evening outdoors. *An. stephensi* and *An. arabiensis* both feed at night, however, *An. stephensi* is mostly endophilic while *An. arabiensis* is exophilic.

In addition to species level differences, the different patterns of *kdr* variation may be explained by multiple evolutionary processes acting on each population sampled: 1) The data may reflect different levels of selective pressure occurring at each location, such that the populations that were under selective pressure from insecticides exhibited *kdr* mutations and no intronic variation. 2) The variation could also reflect previous demographic events, like recent drops in population size or population introductions resulting in a bottleneck and a decline in intronic variation. We can best evaluate these possibilities in the context of variation at other regions of the genomes in these mosquitoes. The COI have been previously analyzed in the *An. arabiensis* and *An. stephensi* (Carter et al. 2018, Carter el 2019). While multiple COI haplotypes were observed for each *An. arabiensis* collection, only a single COI haplotype was identified in the *An. stephensi*. The higher level of diversity in COI in *An. arabiensis* relative to the *kdr* intronic region supports that selective pressure rather than population bottleneck has shaped the variation at the *kdr*. The opposite pattern observed in the *An. stephensi* gives greater support for the absence of selection on that locus in that species. The degree of variation at kdr in *An. stephensi* may also reflect the likely notion of this species being a recent introduction to that region, so it would not have had the same years of exposure to the local pressure that would cause evolved target-site resistance in the local vector populations. No COI data was available for the *Cx. pipiens* s.l. in this study, and both population bottleneck and/or selection on the *kdr* locus remain plausible explanations for the lack of variation.

The multiple collections that comprised our *An. arabiensis* sample set provide preliminary insight into the basis for population *kdr* variation within a species. We observed a range of *kdr* allele frequencies across the *An. arabiensis* sample collections. The collections differ by location and/ or date of collection, suggesting the geography or timing could play a role in the variation in kdr L014 frequencies observed. Additional surveillance in a larger sample size is needed to verify the importance of geographic and temporal factors shaping the frequency of the mutation. Another notable observation, was the shared intron haplotype between the *An. arabiensis* that carried the L1014F mutation to those that did not. The mosquitoes that carried the once advantageous allele may suffer fitness costs in the absence of the selective pressure, which would result in a rebound of the wild-type allele at that locus. These findings underline the value of investigating the *kdr* intronic variation for evidence of fluctuating selective pressures and the potential for the emergence of insecticide resistance in the future.

Several limitations to these studies should be considered. The *An. stephensi* were collected as larvae and pupae and the *An. arabiensis* and *Cx. pipiens s.l.* were collected as wild-caught adults. This method of collection may pose a concern that the immature specimen set would not reflect the natural diversity of the wild-caught adult population. Concerns with clonality however are lowered when considering the level of diversity observed at the *An. stephensi kdr* locus and at the ace-1R locus (3 haplotypes detected; data not shown). In addition, while *An. stephensi* phenotypic resistance has been reported for east Ethiopia, phenotypic data on *An. arabiensis* and its association with *kdr* has only been studied for portions of the country outside of east Ethiopia. Also, the association of *kdr* mutations and phenotypic resistance in *Cx. pipiens sl* observed in other parts of the world have not been confirmed in Ethiopia. Follow-up studies would benefit from additional bioassay tests for *An. arabiensis* and *Cx. pipiens* in east Ethiopia in junction with the molecular analysis of *kdr*. Finally, given the geographic variation in kdr mutation frequencies observed in *An. arabiensis*, future studies should look at the frequency of *kdr* mutations of these vectors in other regions in Ethiopia to confirm the status of target-site pyrethroid/ DDT resistance.

In conclusion, the different patterns of diversity at the *kdr* loci across species indicate that Culicidae in east Ethiopia likely have different mechanisms of resistance profiles. Both *An. arabiensis* and *Cx. pipiens* sample sets revealed notable L1014F allele frequencies that confer target-site resistance and absence of intron variation that tells of selective pressure on that locus in those species. Additional investigations are needed to determine the mechanisms and genetic basis of pyrethroid resistance (metabolic, cuticle, or another undiscovered mechanism) in *An. stephensi*. These finding emphasize the need for careful consideration of molecular approaches used to evaluate insecticide resistance status across multiple species and will inform the development and future implementation of novel integrated vector control strategies.

## Abbreviations

BLAST: Basic Local Alignment Search Tool
DNA: Deoxyribonucleic Acid
FMOH: Federal Ministry of Health
ITS2: Internal transcribed spacer 2 region
NCBI: National Center of Biotechnology Information
PCR: Polymerase chain reaction
*KDR*: knockdown resistance
*VGSC*: voltage-gated sodium channel
COI: Cytochrome c oxidase subunit 1 gene
CDC: Centers for Disease Control and Prevention
WHO: World Health Organization

## Declarations

### Competing interests

The authors declare that they have no competing interests and the manuscript has not been published before or submitted elsewhere for publication.

### Availability of data and material

The datasets supporting the conclusions of this article are included within the article and its supplemental files. Sequences have been submitted to NCBI Genbank database.

### Funding

This study was financially supported by Jigjiga University. This project was also funded by Baylor University and the University of North Carolina at Charlotte Multicultural Postdoctoral Fellowship.

## Acknowledgements

Our gratitude goes to Mr. Negib Abdi and Habtamu Atlaw for their facilitating financial and arranging the car for the field work. We would also like to thank Mr. Geleta Bekele for his technical support in rearing mosquito at field laboratory. We would like to also thank Dr. Jason Pitts and Ms. Jeanne Samake for the helpful comments on the manuscript. We acknowledge the support of various entities of the University of North Carolina at Charlotte including: the Department of Bioinformatics and Genomics of the College of Computing and Informatics and the Department of Biological Sciences of the College of Literature Science and the Arts. We are grateful for the support of the Belk Family. We would also like to acknowledge the support of Baylor University Department of Biology of the College of Arts and Sciences.

## Notes

### Competing Interest Statement

The authors have declared no competing interest.

### Summary of Updates

Figures updated. Methods and results refocused on the genetic data.

## References

1. Organization WH, 2019. World malaria report 2019.

2. Organisation WH, 2018. World malaria report 2018.

3. Balkew M, et al., 2020. Geographical distribution of *Anopheles stephensi* in eastern Ethiopia. Parasit Vectors 13: 35.

4. Carter TE, Yared S, Gebresilassie A, Bonnell V, Damodaran L, Lopez K, Ibrahim M, Mohammed S and Janies D, 2018. First detection of *Anopheles stephensi* Liston, 1901 (Diptera: culicidae) in Ethiopia using molecular and morphological approaches. Acta Trop 188: 180–186.

5. Faulde MK, Rueda LM and Khaireh BA, 2014. First record of the Asian malaria vector *Anopheles stephensi* and its possible role in the resurgence of malaria in Djibouti, Horn of Africa. Acta Trop 139: 39–43.

6. Seyfarth M, Khaireh BA, Abdi AA, Bouh SM and Faulde MK, 2019. Five years following first detection of *Anopheles stephensi* (Diptera: Culicidae) in Djibouti, Horn of Africa: populations established-malaria emerging. Parasitol Res 118: 725–732.

7. Golding N, Wilson AL, Moyes CL, Cano J, Pigott DM, Velayudhan R, Brooker SJ, Smith DL, Hay SI and Lindsay SW, 2015. Integrating vector control across diseases. BMC Med 13: 249.

8. Ranson H and Lissenden N, 2016. Insecticide Resistance in African *Anopheles* Mosquitoes: A Worsening Situation that Needs Urgent Action to Maintain Malaria Control. Trends in Parasitology 32: 187–196.

9. Ranson H, N’Guessan R, Lines J, Moiroux N, Nkuni Z and Corbel V, 2011. Pyrethroid resistance in African anopheline mosquitoes: what are the implications for malaria control? Trends in Parasitology 27: 91–98.

10. Davies TG, Field LM, Usherwood PN and Williamson MS, 2007. A comparative study of voltage-gated sodium channels in the Insecta: implications for pyrethroid resistance in Anopheline and other Neopteran species. Insect Mol Biol 16: 361–75.

11. Scott JG, Yoshimizu MH and Kasai S, 2015. Pyrethroid resistance in *Culex pipiens* mosquitoes. Pestic Biochem Physiol 120: 68–76.

12. Ranson H, Jensen B, Vulule JM, Wang X, Hemingway J and Collins FH, 2000. Identification of a point mutation in the voltage-gated sodium channel gene of Kenyan *Anopheles gambiae* associated with resistance to DDT and pyrethroids. Insect Molecular Biology 9: 491–497.

13. David JP, Ismail HM, Chandor-Proust A and Paine MJ, 2013. Role of cytochrome P450s in insecticide resistance: impact on the control of mosquito-borne diseases and use of insecticides on Earth. Philos Trans R Soc Lond B Biol Sci 368: 20120429.

14. Alemayehu E, et al., 2017. Mapping insecticide resistance and characterization of resistance mechanisms in *Anopheles arabiensis* (Diptera: Culicidae) in Ethiopia. Parasit Vectors 10: 407.

15. Asale A, Getachew Y, Hailesilassie W, Speybroeck N, Duchateau L and Yewhalaw D, 2014. Evaluation of the efficacy of DDT indoor residual spraying and long-lasting insecticidal nets against insecticide resistant populations of *Anopheles arabiensis* Patton (Diptera: Culicidae) from Ethiopia using experimental huts. Parasit Vectors 7: 131.

16. Balkew M, Ibrahim M, Koekemoer LL, Brooke BD, Engers H, Aseffa A, Gebre-Michael T and Elhassen I, 2010. Insecticide resistance in *Anopheles arabiensis* (Diptera: Culicidae) from villages in central, northern and south west Ethiopia and detection of kdr mutation. Parasit Vectors 3: 40.

17. Yewhalaw D, Van Bortel W, Denis L, Coosemans M, Duchateau L and Speybroeck N, 2010. First Evidence of High Knockdown Resistance Frequency in *Anopheles arabiensis* (Diptera: Culicidae) from Ethiopia. American Journal of Tropical Medicine and Hygiene 83: 122–125.

18. Yared S, Gebressielasie A, Damodaran L, Bonnell V, Lopez K, Janies D and Carter T, 2020. Insecticide Resistance in *Anopheles stephensi* in Somali Region, Eastern Ethiopia.

19. Tajima F, 1989. Statistical method for testing the neutral mutation hypothesis by DNA polymorphism. Genetics 123: 585–95.

20. Chang X, et al., 2016. Landscape genetic structure and evolutionary genetics of insecticide resistance gene mutations in *Anopheles sinensis*. Parasit Vectors 9: 228.

21. Weill M, Chandre F, Brengues C, Manguin S, Akogbeto M, Pasteur N, Guillet P and Raymond M, 2000. The kdr mutation occurs in the Mopti form of *Anopheles gambiae* s.s. through introgression. Insect Molecular Biology 9: 451–455.

22. Carter TE, Yared S, Hansel S, Lopez K and Janies D, 2019. Sequence-based identification of *Anopheles* species in eastern Ethiopia. Malar J 18: 135.

23. Madden T, 2002. The BLAST Sequence Analysis Tool. Bethesda, MD

24. Singh OP, Dykes CL, Lather M, Agrawal OP and Adak T, 2011. Knockdown resistance (kdr)- like mutations in the voltage-gated sodium channel of a malaria vector *Anopheles stephensi* and PCR assays for their detection. Malaria journal 10: 59.

25. Verhaeghen K, Van Bortel W, Roelants P, Backeljau T and Coosemans M, 2006. Detection of the East and West African kdr mutation in *Anopheles gambiae* and *Anopheles arabiensis* from Uganda using a new assay based on FRET/Melt Curve analysis. Malar J 5: 16.

26. Chen L, et al., 2010. Molecular ecology of pyrethroid knockdown resistance in *Culex pipiens* pallens mosquitoes. PLoS One 5: e11681.

27. Rozas J, Sanchez-DelBarrio JC, Messeguer X and Rozas R, 2003. DnaSP, DNA polymorphism analyses by the coalescent and other methods. Bioinformatics 19: 2496–7.

28. Stephens M, Smith NJ and Donnelly P, 2001. A new statistical method for haplotype reconstruction from population data. Am J Hum Genet 68: 978–89.

29. Scheet P and Stephens M, 2006. A fast and flexible statistical model for large-scale population genotype data: applications to inferring missing genotypes and haplotypic phase. Am J Hum Genet 78: 629–44.

30. Fu YX, 1997. Statistical tests of neutrality of mutations against population growth, hitchhiking and background selection. Genetics 147: 915–25.

31. Fu YX and Li WH, 1993. Statistical tests of neutrality of mutations. Genetics 133: 693–709.

